# Synergistic biodegradation of polyethylene by experimentally evolved bacterial biofilms

**DOI:** 10.1101/2025.04.01.646683

**Authors:** Shan Li, Lei Su, Jiajia Liu, Jingwen Qiu, Lianbing Lin, Ákos T. Kovács, Yicen Lin

## Abstract

Polyethylene, one of the most widely used synthetic polymers, presents significant environmental challenges due to its resistance to biodegradation. Its surface offers a unique ecological niche for microbial colonization and serves as a primary habitat for degrading microorganisms. Despite the pivotal role microbial communities play in plastic degradation, there has been limited research on constructing stable, interacting microbial consortia. In this study, we explored the potential of evolving bacterial biofilm communities to enhance polyethylene degradation. Through long-term experimental evolution, six microbial populations underwent 40 selection cycles using polyethylene as their sole carbon source. The resulting evolved communities formed robust, multi-species biofilms with enhanced degradation capabilities, outperforming their ancestral populations in biofilm production. *Pseudomonas stutzeri* emerged as the dominant species, orchestrating a synergistic interaction with two other isolates through metabolic division of labor. (Meta)-transcriptomics analysis revealed that *Pseudomonas* primarily contributed to the expression of enzymes involved in microbe-mediated degradation of polyethylene, whereas the other community members were responsible for secreting extracellular polysaccharides, improving biofilm formation. This study highlights the potential of experimentally evolved microbial consortia to synergistically accelerate plastic biodegradation, offering promising strategies for environmental bioremediation.

## Introduction

The durability and short lifespan of plastic products, such as polyethylene (PE), have led to the accumulation of plastic waste in the environment^1^. Once introduced, plastic debris creates a unique, unnatural habitat called the “plastisphere”, where microorganisms colonize and grow as multicellular aggregates within a self-produced matrix of extracellular polymeric substances (EPS), forming biofilms^2^. Bacterial biofilms on plastic debris differ from those in surrounding waters, showing lower diversity but a higher abundance of Proteobacteria^3,4^. Although research on soil-based plastisphere communities is limited, previous studies suggest distinct composition unlike those in the surrounding soil^5^. Furthermore, various plastics and their additives promote bacterial biofilm formation^6–8^, underscoring the complex interactions between plastics and microorganisms when addressing environmental concerns.

Microorganisms are increasingly recognized as a promising solution towards reducing plastic pollution^9^. Recent reviews have explored microbial community composition in microplastic environments and their roles in plastic biodegradation^3,10,11^. Environmental plastic samples often host biofilms that contain significant sources of plastic-degrading bacteria^12^. For instance, study has identified key plastic-degrading microbes by comparing bacterial communities between microplastics and sediments^13^. Application of selective pressure can further enhance these biofilms’ degradation capabilities, leading to communities that surpass naturally occurring ones. Consequently, enrichment methods using plastics as the sole carbon source are commonly employed to isolate plastic-degrading bacteria, such as *Bacillus*^9^, *Pseudomonas*^14^, *Rhodococcus*^12^, or others^15^.

However, replicating the described efficiencies outside laboratory settings with monocultures is challenging, as microbial processes primarily occur in mixed-species biofilms. The complexity of the degradation process can be metabolically overburdening for monocultures in addition to constrains of the later stages of metabolism^16^. In contrast, naturally evolved biofilm communities overcome these limitations through metabolic division of labor, where distinct strains perform complementary functions^17^. Cross-feeding of metabolites between species enhances biodegradation efficiency, and spatial organization within biofilms facilitates these interactions^18,19^. For example, initial plastic degradation often occurs within biofilms, with byproducts further broken down by surrounding bacterial communities, enabling complete plastic mineralization^20^. Thus, bacterial communities demonstrate superior environmental adaptability and degradation efficiency compared to monocultures.

Despite growing recognition of bacterial communities in biodegradation, constructing stable, interacting communities remains a challenge. Several studies focus on enrichment methods, where pollutants serve as the sole carbon source, to construct degrading communites^21,22^. Though effective, these methods often overlook the critical role of spatial organization that develops during a long-term biofilm evolution. Experimental evolution (EE) of bacterial biofilms offers an alternative approach, promoting mutualism through repeated cycles of colonization and re-colonization^23^. In addition to directly degrading plastics, the enhancement of microbial biofilms aids in aggregating microplastics and removing larger particles from the environment. For example, Chan et al. used EE to generate microplastic aggregates, which enhanced biofilm formation of *Pseudomonas aeruginosa* and concentrated microplastics to mitigate pollution^24^. Evolved biofilm populations could rapidly diversify into distinct subpopulations that enhance overall fitness and productivity^25^. Over the past decade, biofilms on plastic beads have provided an excellent model for studying biofilm evolution and diversification under experimental conditions^26,27^.

Building on our previous EE approach, we adapted a system for long-term bacterial evolution on plastic surface^28^. A stable, triple-species consortium isolated from an evolved population demonstrated superior PE degradation efficiency compared to monocultures. This enhanced performance was attributed to its increased maintenance capacity and higher extracellular matrix production of the consortium on PE plastic films, resulting from asymmetric cross-feeding and spatial partitioning of biofilm space. Overall, our findings highlight the potential of experimentally evolved bacterial consortia for enhancing plastic waste bioremediation.

## Materials and Methods

### Establishment of experimental evolution, bacterial strain isolation and characterization

Naturally weathered plastic waste was collected from the sediment of a lakeside environment in Dianchi, Kunming, China (102.7671° E, 24.8185° N). The EE method was modified from a previous study on biofilm evolution, where biofilm cells attached to PE plastic floating in a test tube were incubated at 30°C with shaking at 120 rpm. 16S rRNA gene amplicon sequencing was performed at Majorbio, Shanghai, China. Detailed methods are described in the Supporting Information.

### PE degradation assays and determination of PE physical and chemical changes

SEM, FTIR, and WCA were used to assess PE surface morphology, functional group changes, and surface hydrophobicity. Sterilized PE powder was incubated for 30 days at 30°C, then weight loss and LC-MS analysis was performed to determine degradation products, with all assays repeated three times. Detailed methods are described in the Supporting Information.

### Characterization of biofilm formation and cross-feeding experiments

PE films with biofilm were vortexed in liquid carbon-free basal medium (LCFBM), and bacteria were collected by centrifugation, resuspended, and counted using the plate counting method. Spent medium from cultures grown with PE plastic was sterilized and used in cross-feeding experiments, with bacterial growth measured via OD_600_ and productivity assessed as the fold-gain in growth for a week. Detailed methods are described in the Supporting Information.

### Exopolysaccharide characterization

Exopolysaccharide collection was performed as described previously^28^. Strains were cultivated at 30 °C for PE plastic colonization with shaking at 120 rpm. At certain growth stage, bacteria biomass was suspended in 1 mL of 0.9% NaCl buffer and sonicated (5 × 12 pulses of 1 s at 50% amplitude). Bacterial biomass was separated by centrifugation (10 min at 12000g), and the supernatant was collected. Exopolysaccharide content was quantified using the phenol-sulfuric acid method, with a standard curve constructed from diluted glucose solution^54^.

### 16S rRNA fluorescence in situ hybridization and confocal laser scanning microscopy (FISH-CLSM)

Biofilms of evolved populations were formed on PE plastics and fixed with paraformaldehyde. Labeled oligo probes (Cy5 and 6-FAM) were used for detecting specific bacterial groups via FISH. Hybridization and imaging were conducted using CLSM, with 3D visualizations analyzed in ImageJ software. Detailed methods are described in the Supporting Information.

### Transcriptomic analyses

PE pieces (∼2 × 4 cm) served as a substrate for carbon source, and PE-attached bacteria were used for transcriptomic analysis. Triple-species and Ps monocultures were incubated with PE plastics in LCFBM, and LB medium populations served as controls. RNA-seq libraries were prepared, sequenced, and mapped to the Ps genome, with gene expression quantified using FPKM (Fragments Per Kilobase Million) and TPM (Transcripts Per Kilobase Million). Differentially expressed genes were identified and functionally analyzed using KEGG and GO enrichment tools. Detailed methods are described in the Supporting Information.

## Results

### Parallel evolution of bacterial populations on PE plastics

To identify bacterial communities capable of forming biofilms and degrading polyethylene (PE), we collected plastic debris from contaminated sediment in Dianchi Lake, Yunnan, China. After an enrichment experiment with PE plastics, we established six parallel bacterial cultures and subjected them to experimental evolution (EE) under controlled laboratory conditions. Strains that successfully colonized PE formed multi-species biofilms within 72 hours. These biofilms were transferred to fresh tubes with new medium and PE for re-colonization (Figure 1A). All six evolved populations increased biofilm productivity in the EE regime, suggesting improved PE degradation and possible synergy among the strains (Figure 1B). The evolved communities formed strong biofilms on the PE films, with minimal bacterial presence in the surrounding medium (Figure S1a). Degradation tests confirmed that the evolved communities produced more biofilm on PE than the initial cultures (Figure S1bc).

**Figure 1.**
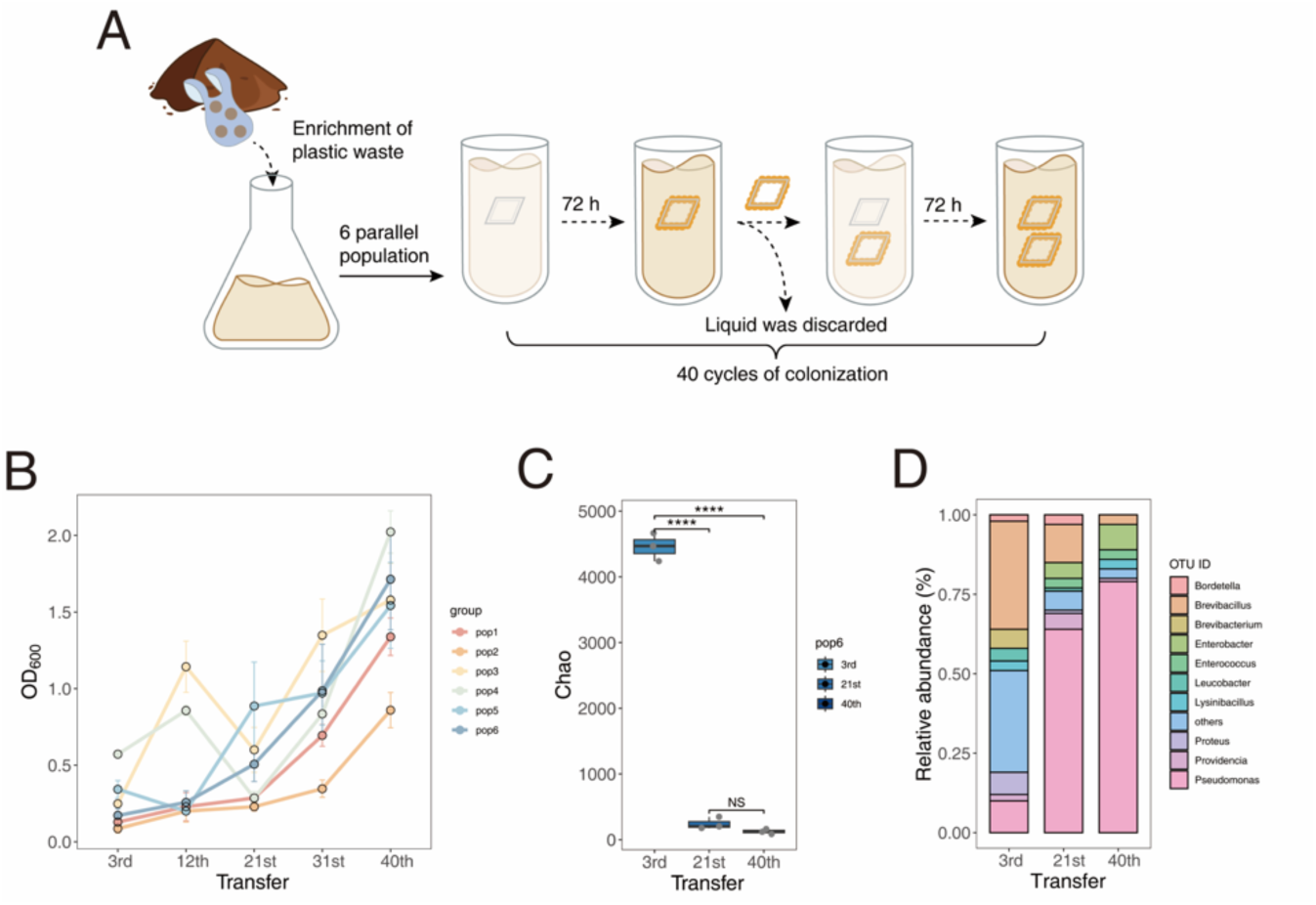
An analysis of the process by which experimental evolution shapes biofilms on PE plastic surfaces. (A) Description of the specific process of experimental evolution, associating with bacterial biofilm formation and dispersal of cells. (B) Quantitative analysis of biofilms on PE plastic surfaces at five stages during evolution. (C) 16S amplicon analysis of pop6 indicated that bacterial alpha diversity within the biofilm rapidly decreased as experimental evolution progressed. (D) Analysis of bacterial community composition at the genus level in biofilms as the experiment progressed. Error bars represent standard error of the mean of three biological replicates (n = 3). \*\*\*\**p* < 0.0001, NS: not significant, based on ANOVA analysis.

To study the evolutionary dynamics of bacterial communities, biofilms were collected at three time points from population 6 (pop6), chosen for its consistently increasing biofilm production. Total DNA was then extracted for 16S rRNA amplicon sequencing. Alpha diversity indices were used to analyze changes in biofilm complexity. Compared to the ancestral populations, the evolved populations showed a sharp decline in community richness and diversity between stages S1 and S2, with no significant changes between stages S2 and S3 for Chao1 (Figure 1C) and Shannon index (Figure S2a). Beta diversity analysis also revealed significant separation in the early stages of evolution (Figure S2b). These results suggested that the experimental conditions created a strong evolutionary bottleneck, driving rapid adaptation to the environment, where hydrocarbons were the sole carbon source. Furthermore, *Pseudomonas* species became dominant during EE, comprising over 60% of the community by the mid-phase, indicating their fitness advantage (Figure 1D).

At the end of the experiment, bacterial biofilms on the PE surface were collected in PBS buffer and vortexed with glass beads to detach the cells from the PE substrate. The resulting bacterial suspension was streaked onto LB agar plates to obtain culturable bacterial isolates. Through 16S rRNA gene amplification and sequencing, three isolates were identified: *Pseudomonas stutzeri* (Ps), *Enterobacter cancerogenus* (Eb), and *Enterococcus casseliflavus* (Ec). After 48 hours of incubation, Ps colonies appeared pale yellow, 4-6 mm in size with serrated edges, likely reflecting their strong biofilm-forming capacity. Eb colonies were larger, semi-transparent with clear edges, whereas Ec colonies were the smallest, about 2 mm in diameter, yellow in color, and smooth (Figure 2A).

**Figure 2.**
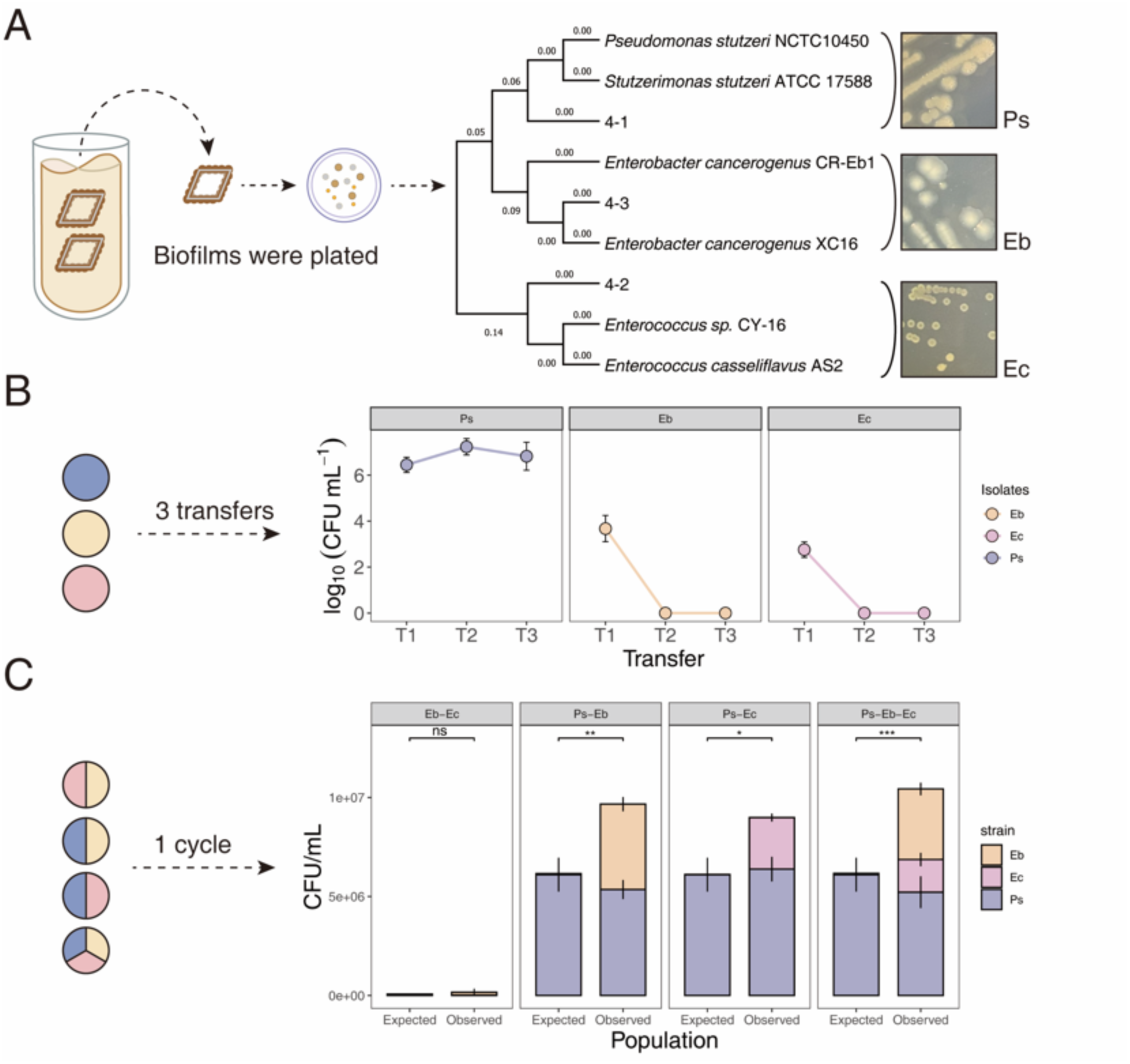
Three dominant bacterial strains were isolated from the group with the highest biofilm production, and their stability was analyzed. (A) The biofilm on the final piece of plastic was vigorously vortexed, plated on LB agar, and the bacteria were isolated and identified. (B) Stability of biofilm formation by isolated strains using PE as the sole carbon source (transferred three times). (C) The expected productivity was calculated by multiplying the initial proportion of each morphotype in the founding population by its monoculture yield (CFU per ml). \**p* < 0.05, \*\**p* < 0.01, \*\*\**p* < 0.001, ns: not significant, based on t-test.

To test the PE colonization and degrading ability of the isolated strains, monocultures were grown with PE plastic as the sole carbon source, allowing biofilm formation. Biofilms were transferred three times following the EE protocol, and bacterial counts were measured as colony-forming units (CFU) per mL. Only Ps consistently utilized PE and formed stable biofilms across all three transfers. In contrast, Eb and Ec formed biofilms during the initial inoculation but showed insufficient biofilm formation after two subsequent transfers on new plastic surfaces (Figure 2B). This inability to persist may be due to their limited capacity to use PE as a carbon source, especially during the re-colonization stages.

To explore potential interactions among these isolates on PE surface, we tested pairwise co-cultures and the three members synthetic community (SynCom) for their ability to utilize PE and form biofilms. Although the co-culture of Eb and Ec led to a slight increase in biofilm production, the difference was not significant compared with the expected yield (Figure 2C). However, when Ps was present, all co-culture combinations showed a significant increase in biofilm production compared with the expected yield based on mono-culture productivity (Figure 2C). This indicated that Ps plays a dominant role in promoting biofilm growth, leading to a more than twofold increase in the total population size, facilitated by potential complementarity. Ps appears to act as the keystone species in these interactions, enhancing the growth of the other strains. We hypothesized that Ps is more efficient at degrading PE, whereas Eb and Ec, likely due to the shorter cultivation periods, cannot effectively utilize the degradation products when cultured individually.

### Evolved bacterial consortia promoted PE plastic degradation

Since PE plastic serves both as an attachment surface and as the sole carbon source, increased biomass likely reflects enhanced degradation. Ps plays a key role in the microbial community due to its consistent colonization of PE. Therefore, we compared the degradation efficiency of Ps mono-culture with the evolved microbial community.

SEM images showed depressions on the PE surface treated with Ps, unlike the smooth control, confirming that Ps can degrade PE (Figure S3a). The SynCom created even more extensive surface damage, indicating greater PE degradation. Hydrophobicity, measured by water contact angle (WCA), decreased from 105.1 ± 1.4 in the control to 101.1 ± 1.0 in Ps-treated and to 96.1 ± 2.2 in SynCom-treated samples, with the latter showing a greater reduction (Figure S3b). The samples treated with microbial community showed a greater reduction in hydrophobicity compared to Ps alone.

Weight loss experiments also confirmed higher PE degradation by the community than by Ps alone (Figure 3A). Fourier Transform Infrared (FTIR) imaging was applied to analyze changes in the surface chemical composition and functional groups of PE plastic (Figure 3B). In the Ps-treated samples, the FTIR spectra revealed two distinct peaks: one at 1650 cm⁻¹ corresponding to C=O stretching, and another at 1120 cm⁻¹ corresponding to C−O stretching. Additionally, in the SynCom-treated samples, a broad peak appeared in the 3000-3600 cm⁻¹ range, corresponding to OH− stretching from carboxylic acids. This suggested that the microbial community-treated PE underwent a higher degree of oxidation and more extensive degradation compared to the Ps-treated samples.

**Figure 3.**
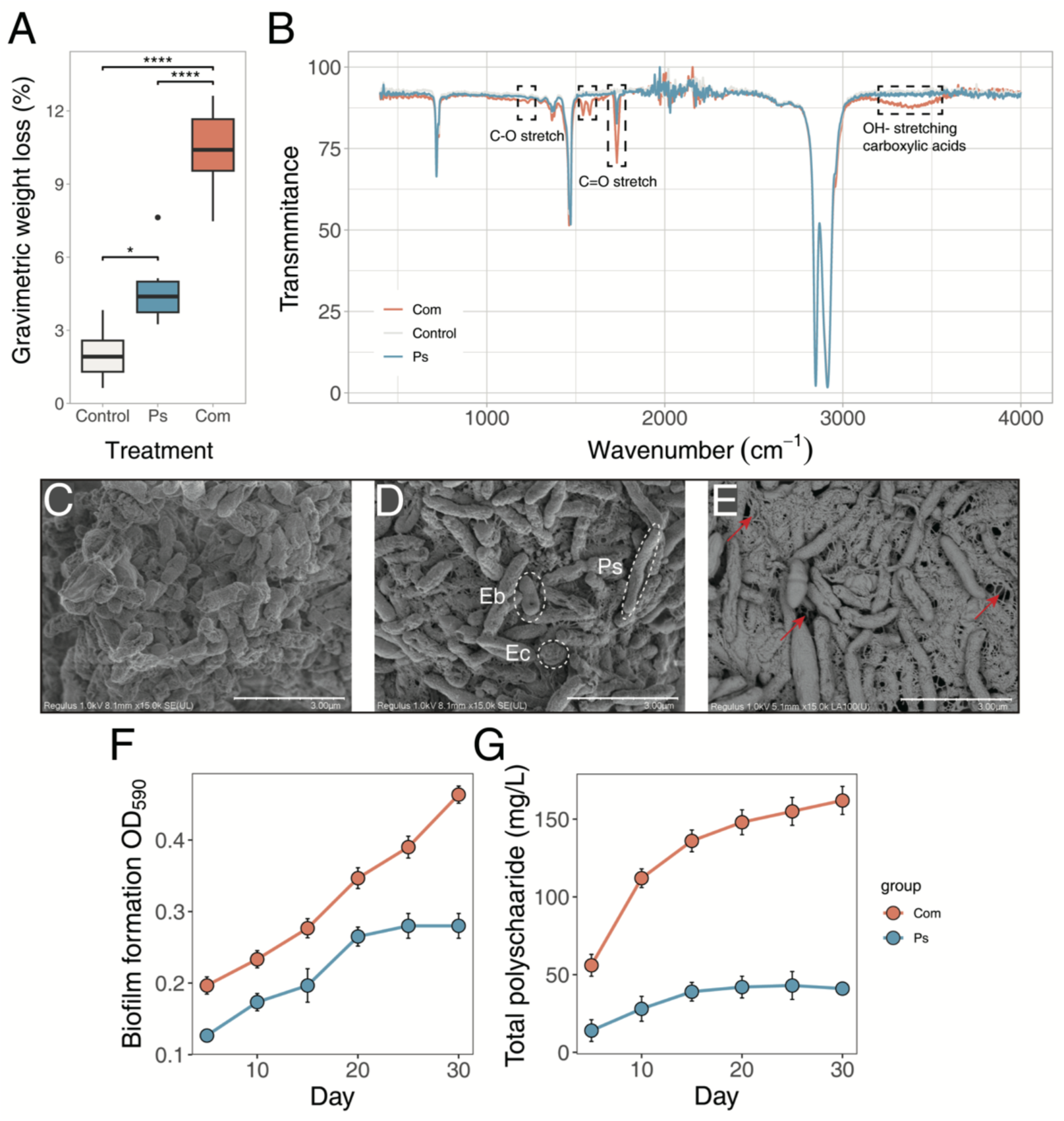
PE degradation and biofilm formation dynamics of Ps and SynCom on the PE surface. (A) Gravimetric weight loss of PE plastic treated with microbial biofilms. (B) FTIR spectrum showed PE functional group changes in the bacteria treated PE samples. (C), (D), (E) Ps, SynCom, and extracellular matrix imaging in SEM, respectively. (F), (G) Time-series analysis of biomass and total extracellular polysaccharides, respectively. \**p* < 0.05, \*\*\*\**p* < 0.0001, based on ANOVA analysis.

We analyzed the degradation products of the PE membrane after 30 days of bacterial culture using HPLC-MS. The results revealed significant differences in the abundance and types of eluted compounds between the two treatment groups and the control group. Differences observed at the end of the mass spectrum strongly suggest that the PE membrane degraded into novel chemical substances (Figure S4). However, due to the lack of standardized reference samples, we were unable to identify the specific chemical characteristics of these degradation products.

### Synergistic biodegradation was achieved by promoted exopolysaccharide secretion of evolved consortium

Both Ps and the SynCom formed microcolonies aiding in plastic degradation (Figures S5a, S5b), but at higher magnification, structural differences were apparent. Ps degraded plastic without a significant extracellular polymeric substance (EPS) network (Figure 3C). In contrast, the SynCom formed a dense biofilm composed of the three strains, including elongated Ps cells, connected by EPS, improving coverage and degradation efficiency (Figures 3D, 3E). Ps likely directed its metabolic resources primarily towards producing degradation-related enzymes rather than extracellular matrix.

We compared the biofilm growth and total polysaccharide production, which reflected EPS production, during colonization. By day five, both Ps and the SynCom showed sufficient bacterial attachment to the PE plastic (Figure 3F). However, the SynCom consistently maintained a higher bacterial abundance. whereas Ps reached a stable phase around day 20 with no further growth, the SynCom continued to increase steadily (Figure 3F). Additionally, total carbohydrate production, a key component of EPS, further explained the SynCom’s stronger persistence on PE plastic. Over the 30-day period, Ps’ carbohydrate production increased gradually from 14 mg/L to about 40 mg/L, whereas carbohydrate production of the SynCom peaked at around 162 mg/L (Figure 3G). This significant difference aligned with the SEM observations, indicating that the metabolic cooperation within the SynCom enhanced EPS secretion and biofilm formation. The mixed-species biofilm structure likely improves molecular transfer efficiency and creates a localized microenvironment that supports enzymatic reactions, which is often a critical bottleneck in plastic degradation.

### Evolved community synergy is attributed to cross-feeding and spatial partitioning

The increased biofilm production and corresponding enhanced degradation efficiency might be the result of two key factors: more efficient nutrient utilization and better use of limited space. Mixed populations were more productive than expected during PE colonization, suggesting that both mechanisms are likely at play (Figure 2C). To quantify the effects of cross-feeding alone, we measured the growth of Ps, Eb, and Ec in the spent medium supernatants from both their own and the other isolates. All isolates - except Ec in its own spent medium - showed enhanced growth in supernatants compared to the original PE medium (Figure 4A). However, the degree of growth enhancement varied, with benefits being asymmetrical. Ps grew best in its own supernatant, increasing its productivity nearly fivefold compared to unconditioned medium, which is expected given its primary role in degrading PE and dominating the evolved population. In contrast, Eb grew optimally in Ps supernatant, whereas Ec thrived in the spent medium of the SynCom. Ec benefited significantly from the metabolic by-products of Ps and Eb, forming an interdependent food web in which only Ps can sustain itself independently (Figure 4B).

**Figure 4.**
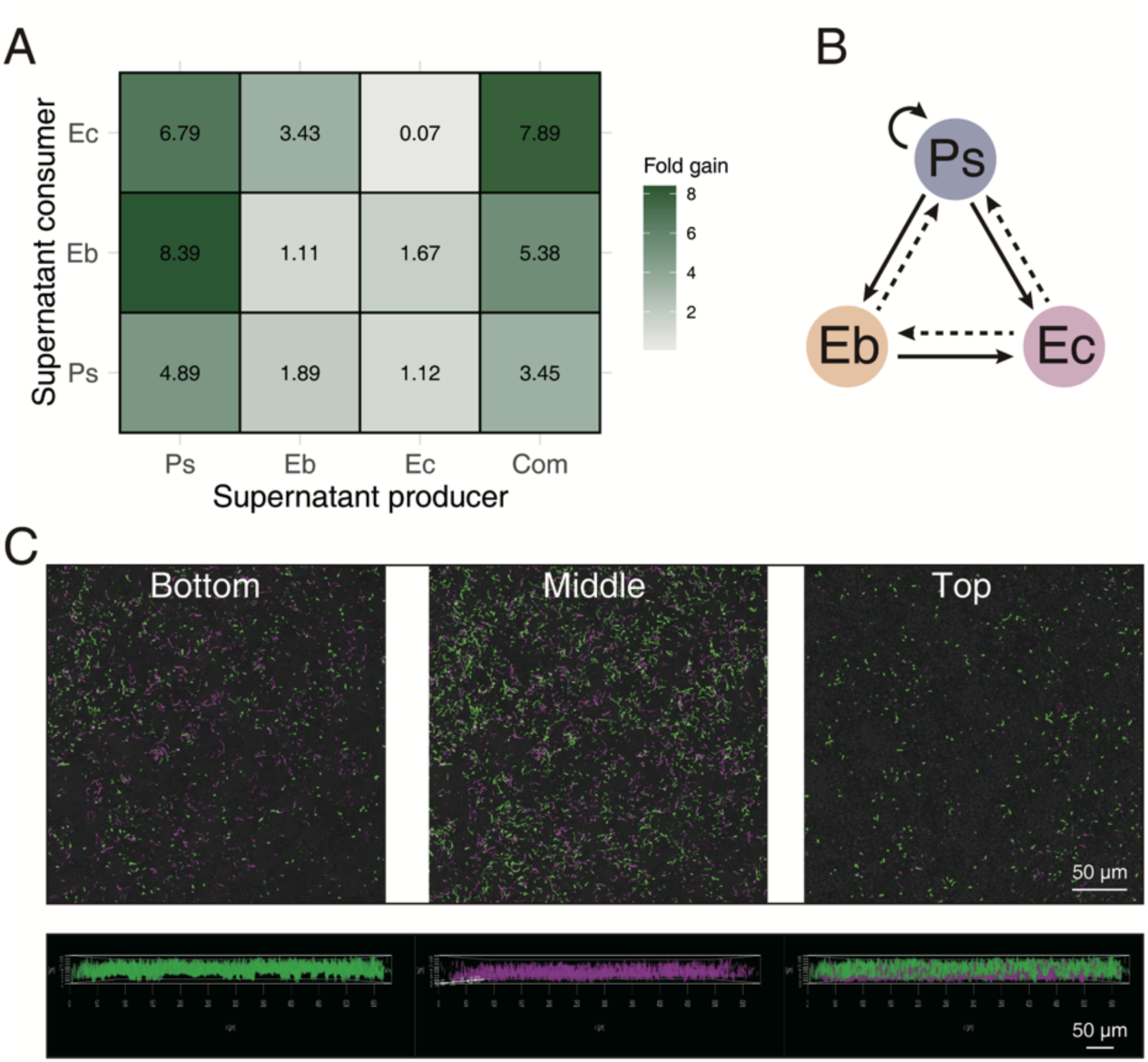
Cross-feeding interactions between isolates and CLSM imaging analysis of biofilm structure. (A) Pair-wise culturing assay with the numerical values represents the fold increase in growth in the supernatant relative to the growth in unconditioned medium. (B) Bacterial interaction diagram: solid lines indicate promotion, whereas dashed lines represent neutral effects (no promotion). (C) CLSM imaging of SynCom with Ps false colored in magenta, whereas the other two strains in green.

To investigate how the isolates partition biofilm space, we used fluorescence in situ hybridization combined with confocal laser scanning microscopy (FISH-CLSM) to image the mixed population. Since Ps and the other two isolates exhibited distinct growth patterns during biofilm assembly, Ps was stained with the cyanine 5 (Cy5, magenta), whereas Eb and Ec were stained with 6-carboxyfluorescein (6-FAM, green). Ps and the other two isolates contributed to the biofilm structure in distinct ways (Figure 4C). Ps adhered more effectively to the plastic surface, spreading horizontally rather than growing vertically. In contrast, Eb and Ec formed the tallest and densest aggregates, covering the biofilm surface. These findings suggested that Ps occupied the basal layer of the biofilm, degrading the PE plastic, whereas Eb and Ec attached to the degradation by-products and formed higher aggregates that encapsulated Ps and the plastic. This structure created an optimal environment for Ps to degrade PE, as it led to a high concentration of enzymes within the biofilm. After disrupting the SynCom structure (biofilms were sonicated and transferred), total biofilm production gradually decreased, eventually leading to biofilm collapse (Figure S6).

### Gene expression of Ps vs. SynCom during PE plastics degradation

Differential gene expression was recorded in Ps alone and the SynCom, and calculated as log2 fold change (LogFC). Principal component analysis using the Bray-Curtis distance matrix was performed to assess expression profile variation. The three replicates of PE-colonizing Ps clustered tightly that demonstrates the reproducibility and consistency of the transcriptomic data (Figure S7a). Differential gene expression analysis using DESeq2 identified 293 genes that were significantly upregulated under PE colonizing conditions compared to the reference nutrient-rich medium (p < 0.05, Table S1). The fold changes of these upregulated genes ranged from 2 to 141. In contrast, 229 genes were downregulated, and 3673 genes showed no significant difference between the two growth conditions (Figure 5A). Genes differentially expressed in Ps cells cultured under PE-colonizing and planktonic lysogenic broth (LB) conditions exhibited two distinct patterns (Figure S7b).

**Figure 5.**
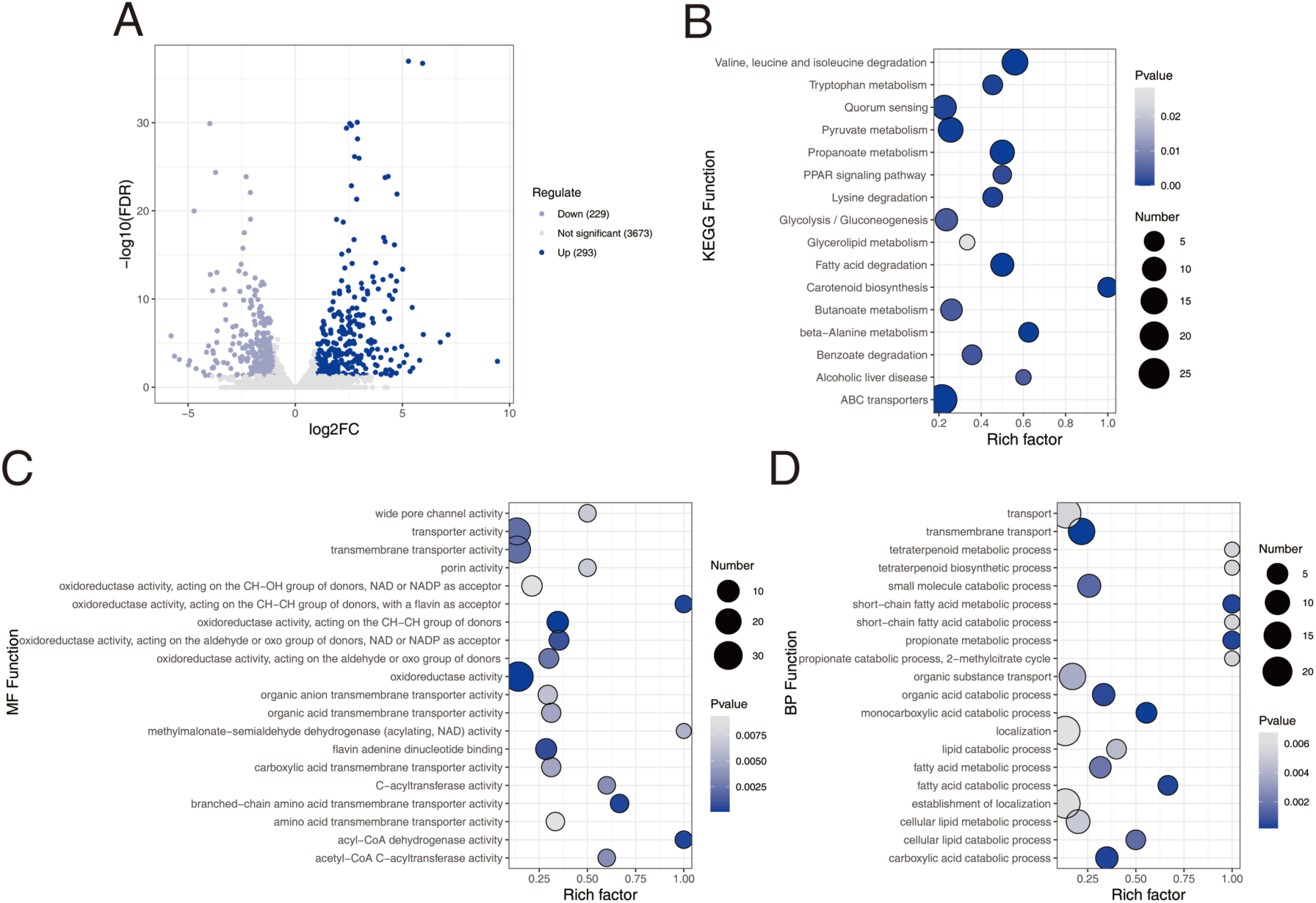
Volcano plots of differentially expressed genes and functional gene enrichment analysis. (A) Volcano plot of differentially expressed genes in Ps during PE plastic degradation compared to growth in LB medium. (B) Enriched analysis of KEGG functions in Ps during PE plastic degradation compared to growth in LB medium. (C) Enriched molecular functions of GO terms in Ps during PE plastic degradation compared to growth in LB medium. (D) Enriched biological process of GO terms in Ps during PE plastic degradation compared to growth in LB medium.

Following annotation, KEGG (Kyoto Encyclopedia of Genes and Genomes database) and GO (Gene Ontology) enrichment analyses were conducted to identify functional changes in cells grown in the presence of PE. Upregulated genes mapped to 16 significantly enriched KEGG pathways. These included pathways related to PE degradation, such as fatty acid metabolism and the breakdown of polyethylene by-products (pyruvate, propanoate, and butanoate) In addition to pathways associated with bacterial activities, such as quorum sensing and macromolecular transport (e.g., ABC transporters) (Figure 5B). The GO enrichment analysis identified 80 enriched GO terms (Table S2). The top 20 enriched molecular functions included transport of by-products, including wide pore channels, transmembrane transporters, and organic and amino acid transport (Figure 5C). Most remaining molecular function pathways related to the oxidation of polyethylene, targeting CH-OH or CH-CH groups, as well as beta-oxidation of polyethylene by-products. Biological processes were strongly enriched in the metabolism of fatty acids, organic acids, monocarboxylic acids, lipids, and carboxylic acids, all involved in the depolymerization and cellular metabolism of polyethylene (Figure 5D). The most enriched biological process pathways related to short-chain fatty acids, propionates, and tetraterpenoids. The *prp* operon, consisting of five genes (*prpB*, *prpC*, *acnD*, *prpF*, and *prpD*) in *Pseudomonas spp.*, is responsible for propionate metabolism, which is a downstream product of odd-chain-length alkane oxidation. All five genes were significantly upregulated in the presence of PE, suggesting a core degradation cycle and energy route producing pyruvate (Figure S8).

We tested the expression patterns of the SynCom community and the function-species correlations during PE utilization. After quality control and assembly with Trinity, differential expression analysis (edgeR, FDR < 0.05, |log2FC| > 2) identified 30485 transcripts, including 9660 upregulated and 4518 downregulated transcripts (Figure 6A). Upregulated genes, categorized via KEGG, were predominantly associated with quorum sensing, biofilm formation, and carbon metabolism (Figure 6B). GO analysis revealed enriched functions in PE-treated samples, particularly in transferase activities and membrane-related processes, underscoring the role of intercellular communication in PE degradation (Figure S9). Species-specific expression patterns showed dominant genes from Eb in LB and Ps in PE (Figure 6C, 6D), highlighting Ps’ contributions to key metabolic pathways in the presence of PE were adaptive.

**Figure 6.**
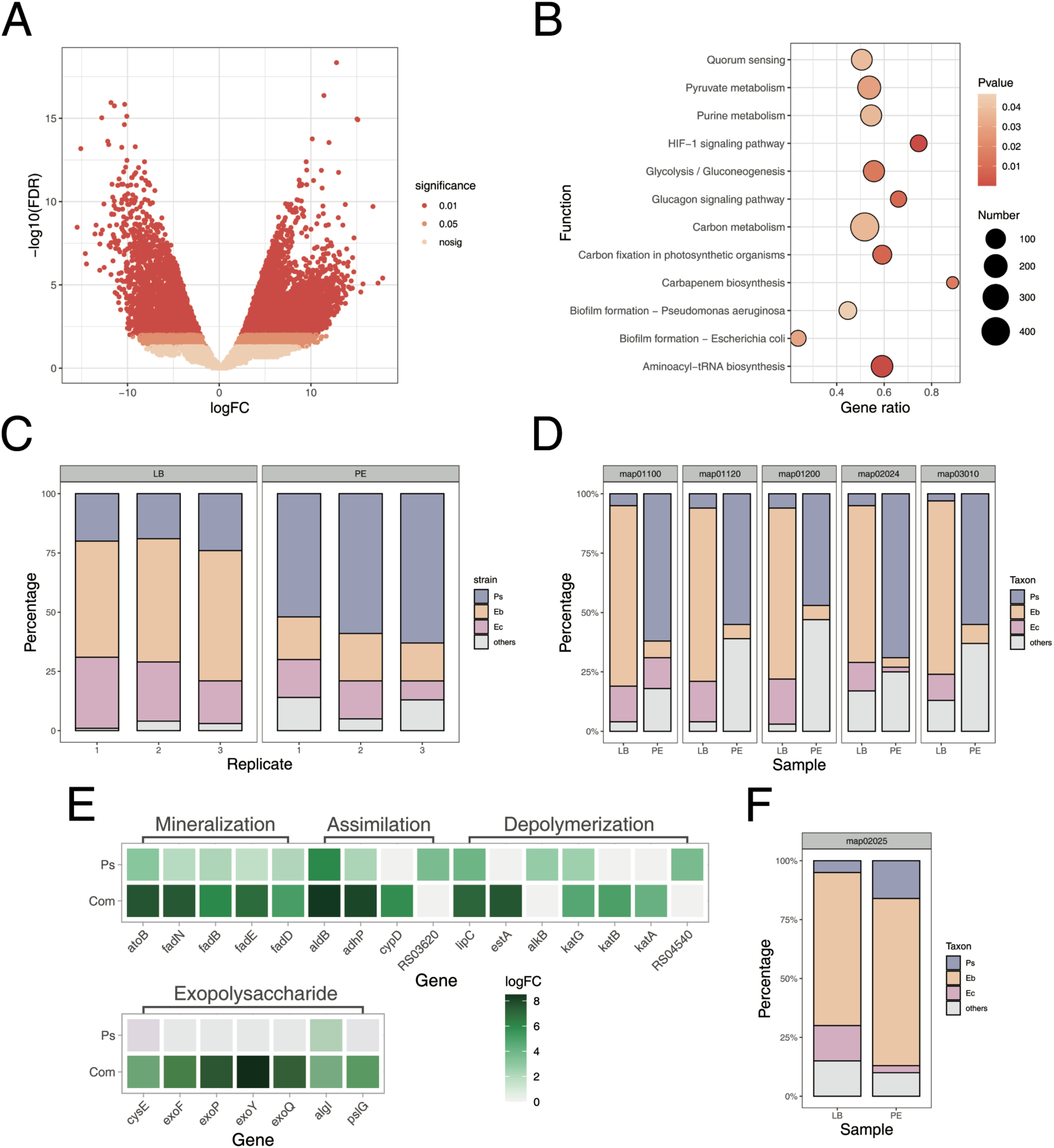
Functional and expressive contribution analysis of metatranscriptomic data. (A) Volcano plot of differentially expressed genes in the SynCom during PE plastic degradation compared to growth in LB medium. (B) Enriched analysis of KEGG functions in Com during PE plastic degradation compared to growth in LB medium. (C) The contribution proportions of three isolates to the total transcripts in the SynCom during PE degradation and LB culture. (D) The contribution proportions of three isolates to the total transcripts in the top five KEGG pathways under two culture conditions. (E) Specific gene expression related to the process of PE degradation and biofilm formation. (F) Relative KEGG pathways of isolates expressive contributions.

To thoroughly compare the degradation processes between monocultures and the bacterial community, the gene expression profiles involved in specific degradation pathways were dissected. During the depolymerization of PE, the *katABG*, *estA*, and *lipC* genes were upregulated in the SynCom, whereas *alkB* was upregulated in Ps monocultures (Figure 6E). SynCom and Ps appeared to rely on different extracellular enzymes to introduce oxygen into the hydrocarbon chain and depolymerize PE into low-molecular-weight polymers. In the subsequent oxidation or assimilation process, different enzymes are involved, with P450 hydroxylase (*cypD*) and monooxygenase (RS03620) being significantly upregulated in Com and Ps, respectively. In the following reaction chain, particularly in the β-oxidation process, the genes involved are significantly upregulated in both the SynCom and Ps. Additionally, given the significant secretion of extracellular matrix in the SynCom compared with the Ps mono-culture, expression of genes related to exopolysaccharide production was also examined. In Ps, only the gene associated with alginate transferase was significantly upregulated, whereas in the SynCom, seven genes related to exopolysaccharide production were upregulated, confirming the results related to increased biofilm matrix production in the SynCom (Figure 6E). In the KEGG function (map02025) related to biofilm formation, Eb played a dominant role rather than Ps, suggesting a task division within the SynCom (Figure 6F).

Based on the above results, a conceptual model and central metabolic pathways for PE degradation by Ps and the SynCom can be proposed (Figure 7). The most significant difference between the degradation processes of monocultures and the bacterial community is the involvement of extracellular enzymes, including catalase-peroxidase KatG, monooxygenases, lipases, and esterases. After the absorption of small-molecule alkanes, Ps and the SynCom relied on monooxygenases and cytochrome P450 enzymes, respectively, to produce alcohol. Importantly, in the later stages of metabolism, key genes were significantly upregulated in both Ps and the SynCom.

**Figure 7.**
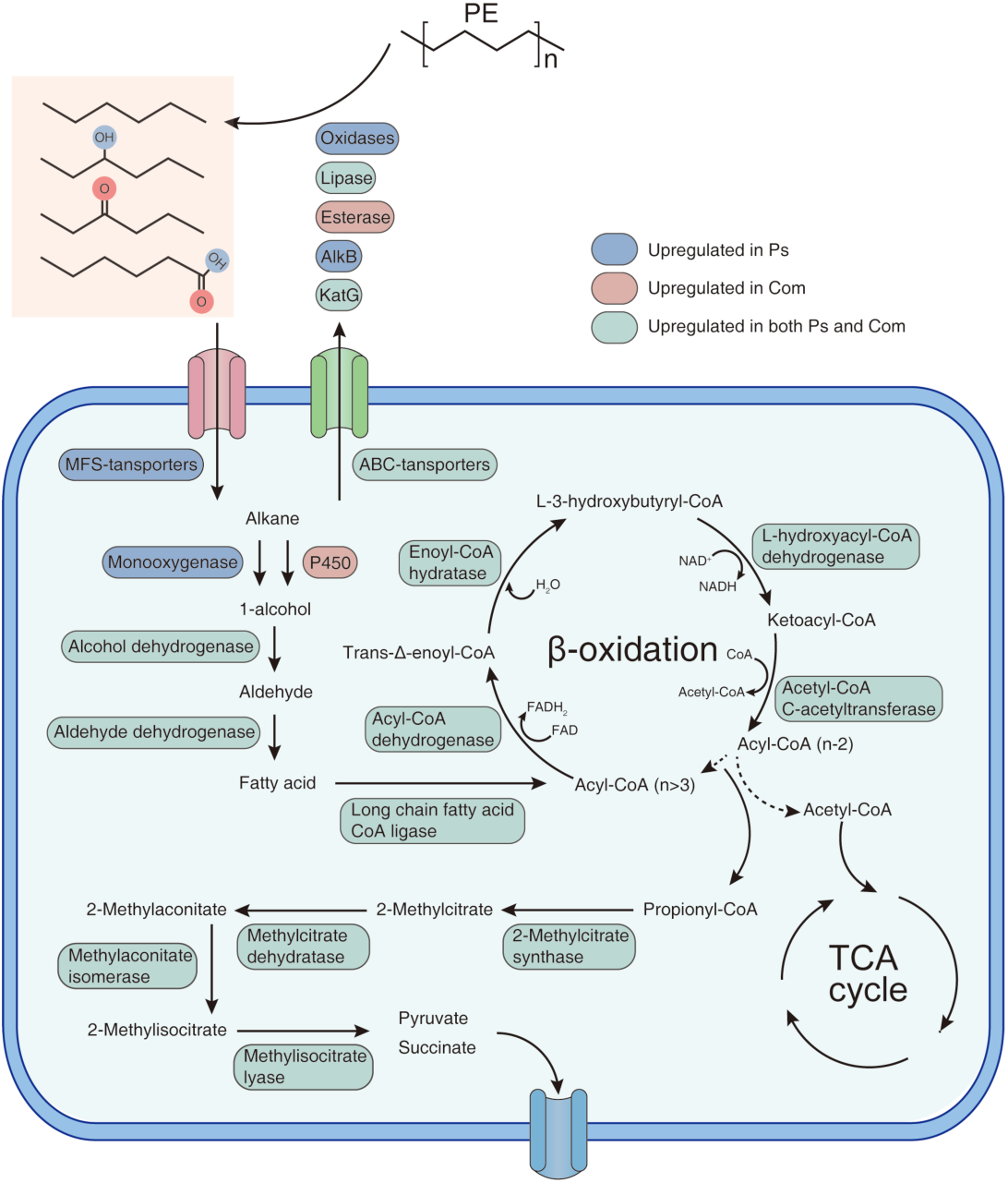
Conceptual model of Ps and the SynCom utilizing PE as the sole carbon source during PE degradation. The blue, red, and green boxes represent features identified only in Ps, only in the SynCom, and in both Ps and the SynCom, respectively.

## Discussion

Bacterial metabolic cooperation in biofilms has been described in various natural and artificial environments, including food^29,30^, plant roots^31^, and laboratory systems^32,33^. However, aside from bioinformatics models^34^, few studies managed to dissect how to create these communities reliably in the laboratory. The primary key finding of our study is that EE effectively creates metabolically cooperative bacterial populations. Microbial plastic degradation is closely linked to environmental plastic pollution, highlighting that microbes promptly adapt to these selection pressures over time^35^. As a result, several studies have used plastic biofilms to isolate plastic-degrading bacteria successfully^36,37^. One key benefit of microbial communities, compared to single strains, is that microbes often exchange metabolic byproducts, which leads to cross-feeding^19,38^. For example, in nitrogen-poor environments, nitrogen-fixing bacteria and degraders can work together to break down pollutants^39^. In marine microbial communities, cross-feeding of metabolites like n-alkanals and aromatics aids bacterioplankton formation^40^. Studies on cross-feeding showed that different species specialize in certain metabolic pathways, like amino acid or sugar synthesis, making them more efficient^41–43^. Our results demonstrated that evolved microbial communities perform superior to individual strains in both survival on PE surfaces and plastic degradation. Metabolic cooperation benefits the community composed of *Pseudomonas*, *Enterobacter*, and *Enterococcus* species (Figure 2, 3). Specifically, *Pseudomonas* and its byproducts in the supernatant help the whole community to grow and therefore act as a keystone species.

Moreover, repeated selection through EE led to a stable formation of structured communities on plastic surfaces, suggesting that EE overcame the main challenge of enriched communities: their initial detriment to form structured populations. CLSM imaging revealed distinct spatial distribution of Ps, Eb, and Ec within the biofilm, suggesting an ecological task division similar to differentiating ecotypes of experimentally evolving biofilms^25,26,44^. Ps initiates the degradation of PE plastic and is predominantly localized at the base of the biofilm, near the plastic substrate. In contrast, Eb and Ec form the protective outer layers that potentially enhance the biofilm’s stability and thickness. Moreover, both biomass and extracellular matrix production are significantly higher in the SynCom compared with the Ps mono-culture. Increased biofilms could potentially concentrate the enzyme locally around the target substrate and retain it within the biofilm matrix for extended periods, thereby enhancing the overall rate of plastic degradation^45^. Such cooperative interaction not only maximizes the biofilm’s structural resilience but also creates a microenvironment that enhances the degradation process, which could be leveraged in future bioremediation strategies. Similarly, a previous study reported that biofilm culture methods are more effective than enrichment methods for studying community-level interactions in PAH-degrading microbial communities, as biofilm culture better preserved active bacterial and genetic diversity^46^. This is attributed to a major concern of enrichment assays: the emergence of so called “cheaters” that exploit benefits produced by the “cooperators”. In well-mixed, unstructured environments cooperators are more susceptible to exploitation by cheaters^47^. In the described SynCom, *Pseudomonas* act as a public producer of PE degrading enzymes, whereas Eb and Ec can be considered as either commensals or exploiters. Our results further demonstrated that when the biofilm structure is disrupted, the cooperative species diminishes, potentially leading to biofilm collapse. Cross-feeding in a spatially structured environment can increase antibiotic persistence in both mutualistic *E. coli* and *S. enterica* compared to their respective monocultures^48^. Therefore, establishing a structured microbial community that maximizes cooperation is essential to ensuring efficient PE degradation as demonstrated in our work.

Finally, we dissected the molecular mechanisms through analyszing the gene expression networks of Ps and SynCom under PE induction. Although several studies have demonstrated the PE-degrading capability of *Pseudomonas sp*.^49^, Our study reveals the key pathways and specific genes involved in *Pseudomonas’* degradation capability of hydrocarbon-based organics. Ps upregulated expression of genes related to pathways for alkane degradation and β-oxidation, particularly in the metabolism of even- and odd-numbered carbon fatty acids. Similar to a previous study^50^, the upregulation of the *prp* operon underscored the importance of odd-numbered fatty acid metabolism in PE plastic degradation that indicates a complete degradation of saturated fatty acids.

Metatranscriptomics facilitates the study of multispecies biofilm communities, particularly in laboratory settings^51^. Analysis of the transcript-to-species ratio indicated that Eb transcription dominated in LB medium, whereas Ps dominated during PE degradation, suggesting that metabolic cooperation is evolutionarily shaped and specifically observable on the PE surface. This applied to the overall transcripts as well as the most common KEGG pathways in the cells, such as map01100 (Metabolic pathways). Quorum sensing (map02024) also showed functional shifts in the strains across the used cultivation media. LB disrupted both community structure and nutrient composition, which are the two most critical factors shaping community cooperation^52^. Comparison of specific genes involved in the degradation process between the SynCom and Ps revealed that the upregulation of genes related to PE degradation was higher in the SynCom than in the Ps mono-cultures, correlating with higher biomass of the SynCom. Regarding the key gene pathways for PE degradation, Ps and the SynCom showed identical pathways after assimilation and mineralization, consistent with previous reports^50,53^. Nevertheless, we observed differential expression of key enzymes, including oxidases, esterases, cytochrome P450, and monooxygenases, during the polymerization process in monocultures and microbial communities, suggesting that the presence of other species influenced the enzyme secretion of Ps. Understanding why these shifts occur will require further analysis, incorporating additional data such as metaproteomics. Through analyzing exopolysaccharide gene and transcript distribution, we found that exopolysaccharide production is primarily enriched in Eb rather than Ps, providing molecular evidence of task division within the SynCom.

In summary, we experimentally evolved natural biofilm populations on PE plastics to describe the molecular mechanisms of community functioning during PE degradation, which has important implications for building artificial microbial communities. The metabolic cooperation within these communities, leading to functional specialization, improves biofilm formation and plays a key role in refining conditions for optimal plastic degradation. Additionally, we identified several extracellular proteins that respond specifically to PE degradation in the evolved populations, suggesting the formation of an enzyme mix (PEases) that could efficiently break down PE.

## Data availability statement

The 16S amplicon sequencing data has been deposited in the NCBI database under accession number: PRJNA1154807. The transcriptomics data of monocultures and cocultures have been deposited in the NCBI database under accession number: PRJNA1153981 and PRJNA1154404.

## Author contribution

S.L. and L.S. conducted the experiments under the guidance of Y.L. A.T.K. and Y.L. conceived the idea and designed the project. All authors contributed to data interpretation, and S.L. and Y.L. wrote the manuscript. All authors reviewed and approved the final version of the manuscript.

## Competing interests

The authors declare no competing interests.

## Acknowledgements

This work was supported by the Yunnan Province Basic Research Program (202401CF070116).

## Notes

### Competing Interest Statement

The authors have declared no competing interest.

## References

1. Roy, P. K., Hakkarainen, M., Varma, I. K. & Albertsson, A.-C. Degradable Polyethylene: Fantasy or Reality. Environ. Sci. Technol. 45, 4217–4227 (2011).

2. Zettler, E. R., Mincer, T. J. & Amaral-Zettler, L. A. Life in the “plastisphere”: microbial communities on plastic marine debris. Environ. Sci. Technol. 47, 7137–7146 (2013).

3. Amaral-Zettler, L. A., Zettler, E. R. & Mincer, T. J. Ecology of the plastisphere. Nat Rev Microbiol 18, 139–151 (2020).

4. McCormick, A., Hoellein, T. J., Mason, S. A., Schluep, J. & Kelly, J. J. Microplastic is an Abundant and Distinct Microbial Habitat in an Urban River. Environ. Sci. Technol. 48, 11863– 11871 (2014).

5. Microbial communities on biodegradable plastics under different fertilization practices in farmland soil microcosms. Sci. Total Environ. 809, 152184 (2022).

6. Wang, H. et al. Nano-sized polystyrene and magnetite collectively promote biofilm stability and resistance due to enhanced oxidative stress response. J. Hazard. Mater. 476, 134974 (2024).

7. Yang, Y. & Alvarez, P. J. J. Sublethal Concentrations of Silver Nanoparticles Stimulate Biofilm Development. Environ. Sci. Technol. Lett. 2, 221–226 (2015).

8. Wang, H. et al. Phthalate Esters Released from Plastics Promote Biofilm Formation and Chlorine Resistance. Environ. Sci. Technol. 56, 1081–1090 (2022).

9. Yang, J., Yang, Y., Wu, W.-M., Zhao, J. & Jiang, L. Evidence of polyethylene biodegradation by bacterial strains from the guts of plastic-eating waxworms. Environ. Sci. Technol. 48, 13776–13784 (2014).

10. Roager, L. & Sonnenschein, E. C. Bacterial Candidates for Colonization and Degradation of Marine Plastic Debris. Environ. Sci. Technol. 53, 11636–11643 (2019).

11. Oberbeckmann, S. & Labrenz, M. Marine Microbial Assemblages on Microplastics: Diversity, Adaptation, and Role in Degradation. Annual Review of Marine Science 12, 209–232 (2020).

12. Tao, X. et al. Polyethylene degradation by a *Rhodococcous* strain isolated from naturally weathered plastic waste enrichment. Environ. Sci. Technol. 57, 13901–13911 (2023).

13. Niu, L. et al. New insights into the vertical distribution and microbial degradation of microplastics in urban river sediments. Water Research 188, 116449 (2021).

14. Howard, S. A. et al. Enrichment of native plastic-associated biofilm communities to enhance polyester degrading activity. Environmental Microbiology 25, 2698–2718 (2023).

15. Yoshida, S. et al. A bacterium that degrades and assimilates poly(ethylene terephthalate). Science 351, 1196–1199 (2016).

16. Wang, C., Huang, Y., Zhang, Z., Hao, H. & Wang, H. Absence of the nahG-like gene caused the syntrophic interaction between Marinobacter and other microbes in PAH-degrading process. J Hazard Mater 384, 121387 (2020).

17. Christensen, B. B., Haagensen, J. A. J., Heydorn, A. & Molin, S. Metabolic Commensalism and Competition in a Two-Species Microbial Consortium. Applied and Environmental Microbiology 68, 2495–2502 (2002).

18. Emadian, S. M., Onay, T. T. & Demirel, B. Biodegradation of bioplastics in natural environments. Waste Manag 59, 526–536 (2017).

19. Xu, X. et al. Modeling microbial communities from atrazine contaminated soils promotes the development of biostimulation solutions. ISME J 13, 494–508 (2019).

20. Meyer-Cifuentes, I. E. et al. Synergistic biodegradation of aromatic-aliphatic copolyester plastic by a marine microbial consortium. Nat Commun 11, 5790 (2020).

21. Gao, R. & Sun, C. A marine bacterial community capable of degrading poly(ethylene terephthalate) and polyethylene. Journal of Hazardous Materials 416, 125928 (2021).

22. Wang, P. et al. Polyethylene mulching film degrading bacteria within the plastisphere: Co-culture of plastic degrading strains screened by bacterial community succession. Journal of Hazardous Materials 442, 130045 (2023).

23. Røder, H. L. et al. Enhanced bacterial mutualism through an evolved biofilm phenotype. ISME J 12, 2608–2618 (2018).

24. Chan, S. Y., Wong, M. W.-T., Kwan, B. T. C., Fang, J. K.-H. & Chua, S. L. Microbial– Enzymatic Combinatorial Approach to Capture and Release Microplastics. Environ. Sci. Technol. Lett. 9, 975–982 (2022).

25. Traverse, C. C., Mayo-Smith, L. M., Poltak, S. R. & Cooper, V. S. Tangled bank of experimentally evolved *Burkholderia* biofilms reflects selection during chronic infections. Proceedings of the National Academy of Sciences of the United States of America 110, E250–259 (2013).

26. Poltak, S. R. & Cooper, V. S. Ecological succession in long-term experimentally evolved biofilms produces synergistic communities. The ISME Journal 5, 369–378 (2011).

27. Mhatre, E. et al. One gene, multiple ecological strategies: A biofilm regulator is a capacitor for sustainable diversity. Proceedings of the National Academy of Sciences 117, 21647–21657 (2020).

28. Lin, Y., Xu, X., Maróti, G., Strube, M. L. & Kovács, Á. T. Adaptation and phenotypic diversification of Bacillus thuringiensis biofilm are accompanied by fuzzy spreader morphotypes. npj Biofilms and Microbiomes 8, (2022).

29. Røder, H. L. et al. Interspecies interactions result in enhanced biofilm formation by co-cultures of bacteria isolated from a food processing environment. Food Microbiology 51, 18–24 (2015).

30. Sadiq, F. A. et al. Dynamic social interactions and keystone species shape the diversity and stability of mixed-species biofilms – an example from dairy isolates. ISME COMMUN. 3, 1– 12 (2023).

31. Yang, N. et al. Interspecific interactions facilitate keystone species in a multispecies biofilm that promotes plant growth. The ISME Journal 18, wrae012 (2024).

32. Liu, W. et al. Low-abundant species facilitates specific spatial organization that promotes multispecies biofilm formation. Environmental Microbiology 19, 2893–2905 (2017).

33. Liu, W., Russel, J., Burmølle, M., Sørensen, S. J. & Madsen, J. S. Micro-scale intermixing: a requisite for stable and synergistic co-establishment in a four-species biofilm. The ISME Journal 12, 1940–1951 (2018).

34. Ruan, Z. et al. Engineering natural microbiomes toward enhanced bioremediation by microbiome modeling. Nat Commun 15, 4694 (2024).

35. Zrimec, J., Kokina, M., Jonasson, S., Zorrilla, F. & Zelezniak, A. Plastic-Degrading Potential across the Global Microbiome Correlates with Recent Pollution Trends. mBio 12, e0215521 (2021).

36. Wang, P. et al. Does bacterial community succession within the polyethylene mulching film plastisphere drive biodegradation? Science of The Total Environment 824, 153884 (2022).

37. Gregory, G. J. et al. Polyethylene Valorization by Combined Chemical Catalysis with Bioconversion by Plastic-Enriched Microbial Consortia. ACS Sustainable Chem. Eng. 11, 3494–3505 (2023).

38. Liu, Z. et al. Metabolite Cross-Feeding between Rhodococcus ruber YYL and Bacillus cereus MLY1 in the Biodegradation of Tetrahydrofuran under pH Stress. Appl Environ Microbiol 85, e01196–19 (2019).

39. Wang, X. et al. Nitrogen transfer and cross-feeding between *Azotobacter chroococcum* and *Paracoccus aminovorans* promotes pyrene degradation. ISME J 17, 2169–2181 (2023).

40. Giordano, N. et al. Genome-scale community modelling reveals conserved metabolic cross-feedings in epipelagic bacterioplankton communities. Nat Commun 15, 1–15 (2024).

41. Tsoi, R. et al. Metabolic division of labor in microbial systems. Proc Natl Acad Sci U S A 115, 2526–2531 (2018).

42. Zhou, K., Qiao, K., Edgar, S. & Stephanopoulos, G. Distributing a metabolic pathway among a microbial consortium enhances production of natural products. Nat Biotechnol 33, 377–383 (2015).

43. Chen, G. et al. Dual Carbon-Chlorine Isotope Analysis Indicates Distinct Anaerobic Dichloromethane Degradation Pathways in Two Members of Peptococcaceae. Environ Sci Technol 52, 8607–8616 (2018).

44. Dragoš, A. et al. Evolution of exploitative interactions during diversification in *Bacillus subtilis* biofilms. FEMS microbiology ecology 93, fix155 (2017).

45. Howard, S. A. & McCarthy, R. R. Modulating biofilm can potentiate activity of novel plastic-degrading enzymes. npj Biofilms Microbiomes 9, 1–10 (2023).

46. Stach, J. E. M. & Burns, R. G. Enrichment versus biofilm culture: a functional and phylogenetic comparison of polycyclic aromatic hydrocarbon-degrading microbial communities. Environmental Microbiology 4, 169–182 (2002).

47. Vetsigian, K., Jajoo, R. & Kishony, R. Structure and Evolution of Streptomyces Interaction Networks in Soil and In Silico. PLOS Biology 9, e1001184 (2011).

48. Xiong, X., Othmer, H. G. & Harcombe, W. R. Emergent antibiotic persistence in a spatially structured synthetic microbial mutualism. The ISME Journal 18, wrae075 (2024).

49. Zhai, X., Zhang, X.-H. & Yu, M. Microbial colonization and degradation of marine microplastics in the plastisphere: A review. Frontiers in Microbiology 14, (2023).

50. Gravouil, K. et al. Transcriptomics and lipidomics of the environmental strain *Rhodococcus ruber* point out consumption pathways and potential metabolic bottlenecks for polyethylene degradation. Environ. Sci. Technol. 51, 5172–5181 (2017).

51. Tan, C. H., Lee, K. W. K., Burmølle, M., Kjelleberg, S. & Rice, S. A. All together now: experimental multispecies biofilm model systems. Environ. Microbiol. 19, 42–53 (2017).

52. Nadell, C. D., Drescher, K. & Foster, K. R. Spatial structure, cooperation and competition in biofilms. Nature Reviews Microbiology 14, 589–600 (2016).

53. Zhang, Y., Pedersen, J. N., Eser, B. E. & Guo, Z. Biodegradation of polyethylene and polystyrene: From microbial deterioration to enzyme discovery. Biotechnology Advances 60, 107991 (2022).

54. Masuko, T. et al. Carbohydrate analysis by a phenol–sulfuric acid method in microplate format. Analytical Biochemistry 339, 69–72 (2005).

