## Supplementary file for "Synergistic biodegradation of polyethylene by experimentally evolved bacterial biofilms"

**Supplementary Texts**

**Supplementary Materials and Methods**

**Text S1. Establishment of experimental evolution, bacterial strain isolation and characterization.** Naturally weathered plastic waste was collected from the sediment of a lakeside environment in Dianchi, Kunming, Yunnan Province, China (102.7671° E, 24.8185° N). Plastic pieces (~2 × 4 cm) were prepared using sterile scissors and incubated in 100 mL of liquid carbon-free basal medium (LCFBM) supplemented with 0.01% yeast extract. The incubation was carried out in 300 mL flasks at 30 °C, with shaking at 120 rpm, for 30 days. Following incubation, the cultures were vortexed and centrifuged to harvest the cells. The collected cells were resuspended in fresh LCFBM as starting cultures for further evolution experiment. LCFBM was prepared with deionized water, containing (per 1000 mL): 0.7 g $KH_2PO_4$, 0.7 g $K_2HPO_4$, 0.7 g $MgSO_4·7H_2O$, 1.0 g $NH_4NO_3$ 0.005 g NaCl, 0.002 g $FeSO_4$, 0.002 g $ZnSO_4$, and 0.001 g $MnSO_4$. Linear low-density polyethylene (LDPE) film (951-050, 30 μm thick) was obtained from SINOPEC Guangdong Maoming Company, China. According to the manufacturer's specifications, this PE product contained no added catalysts or additives.

One milliliter of enriched culture (optical density at 600nm ($OD_{600}$) of 0.1) was inoculated into 14 mL of LCFBM. The culture was incubated in test tubes at 30 °C with shaking at 120 rpm, allowing biofilm formation on floating polyethylene (PE) plastic. In each cycle, a new PE plastic was colonized, and a new biofilm was formed, allowing for a regular selection cycle of attachment and dispersal. The biofilm populations were serially transferred every 72 hours for a total of 40 cycles. Cells that adhered to the tube walls or remained in the planktonic phase were not transferred. Throughout the EE process, biofilm biomass on the PE plastic was monitored using $OD_{600}$ measurements of bacterial suspension, taken five times during the 40 cycles ($3^{rd}$, $12^{th}$, $21^{st}$, $31^{st}$,

$40^{th}$). At the end of the EE, PE plastics with visible biofilms were transferred to disposable culture tubes containing 5 ml of LCFBM with 0.01% yeast extract to promote microbial growth. The culture was vortexed, diluted 1:100 in LCFBM, and 50 μl was plated onto 1:2 diluted LB plates for bacterial strain isolation. Single colonies were streaked twice on LB plates to obtain pure cultures. Overnight cell cultures were then used for Gram staining, following standard protocols.

For both pure cultures and evolved populations, total genomic DNA was extracted using the OMEGA Soil DNA Kit (M5635-02) (Omega, USA), following the manufacturer's instructions, and stored at -20 °C prior to further analysis. The 16S rRNA genes were amplified using primers 27F and 1492R, following standard protocols. Sanger sequencing of the purified PCR products was performed at Sangon Biotech China and further sequences were blasted against NCBI database for strain characterization.

For high-throughput sequencing, 16S rRNA gene V3–V4 region was amplified using 338F (5'-ACTCCTACGGGAGGCAGCA-3') and 806R (5'-GGACTACHVGGGTWTCTAAT-3'). PCR amplicons were purified using VAHTS™ DNA Clean Beads (Vazyme, China) and quantified with the Quant-iT PicoGreen dsDNA Assay Kit (Invitrogen, USA). After quantification, amplicons were pooled in equal amounts, and paired-end 2 × 250 bp sequencing was performed on the Illumina NovaSeq platform using the NovaSeq 6000 SP Reagent Kit (500 cycles) at Majorbio, Shanghai.

**Text S2. PE degradation assays and determination of PE physical and chemical changes.** For the PE powder degradation assay, 0.5 g of PE powder (Sigma-Aldrich, Cat:427772-250g, average $M_n$ ~ 1700 and $M_w$ ~ 4000) was accurately weighed using a fine scale and transferred into a 50 ml centrifuge tube. To sterilize, 20 ml of 70% ethanol was added to the tube, which was then rotated for 30 minutes at room temperature. The sterilized PE powder was collected by centrifugation at 12,000×g for 30 minutes with slow acceleration and deceleration. After removing the supernatant, the PE powder was air-dried overnight under a biosafety hood. All groups were incubated at 30°C for 30 days with shaking at 120 rpm.

Fourier Transform Infrared Spectroscopy (FTIR) was employed to identify the functional group changes over time. A total of 32 scans was captured using smart endurance single bounce with a diamond tip in the 4000 - 400 $cm^{-1}$ range. Spectra were analyzed using Spectragryph software (version 1.2.16). Water contact angle (WCA) analysis was used for analyzing surface hydrophobicity of bacteria-treated and -untreated PE plastics. The contact angle of the plastic films with water was measured at room temperature with the measuring device (HE-CA200). The assays were repeated three times.

The surfaces of the PE plastics, along with the structure of the attached biofilm and microbial community, were examined using a scanning electron microscope (SEM). Both virgin PE samples and bacteria-treated samples collected after 30 days were coated with a 20 nm layer of gold using an SCD 050 sputter coater (Bal-Tec). Further SEM was conducted with prepared samples (Zeiss Sigma 360) and three samples were used for each treatment. After one month of degradation, supernatant was collected and filtered through a 0.2 µm filter. A 2 µL aliquot was detected by LC-MS, equipped with an ACQUITY UPLC BEH C18 analytical column (2.1 × 50 mm, 1.7 µm), at an m/z range of 50–1500[1].

**Text S3. Characterization of biofilm formation and cross-feeding experiments.** The PE films with biofilm were placed in a 50 mL centrifuge tube containing 40 mL of LCFBM and vortexed for 5 minutes, then bacterial suspension was centrifugated for 10 min at 10000 rpm. Pelleted cells were resuspended and counted using the series dilution method of plate counting.

Spent medium were used for cross-feeding experiments. The species were grown in 10 mL LCFBM with commercial PE plastic at pH 7 for 2 weeks at 30 °C. The cultures were centrifuged at 3220×g for 10 min, and the spent medium was sterilized using a 50 mL Steriflip filter unit (0.22 μm; Millipore Sigma). To verify sterility, 50 μL of the spent medium was spotted onto LB plates. The spent medium was used in place of water in the culture medium. A second preculture was prepared for each species, centrifuged, and inoculated at $10^6$ CFU/mL for a one-week incubation. $OD_{600}$ was measured in different spent media, and productivity was defined as the fold-gain in growth (area under the curve of OD600 over 24h), with ratios calculated between two means (n=3) from simultaneous growth assays.

**Text S4. 16S rRNA fluorescence in situ hybridization and confocal laser scanning microscopy (FISH-CLSM).** Biofilms of evolved populations were formed on PE plastics for 2 weeks to ensure bacterial persistence and activity. Once visible bacterial aggregates formed, the PE plastics were dehydrated by placing them on filter paper and soaking in 75% ethanol for 10 minutes at room temperature. Labeled oligo probes, Cyanine 5 (Cy5) and 6-car-boxyfluorescein (6-FAM), were synthesized (Sangon Biotech, China) and used for the detection of *Pseudomonas spp.* [2] (Cy5-labeled probes: 5'-GAT CCG GAC TAC GAT CGG TTT-3'), Enterobacteriaceae[3] (6-FAM-labeled probes: 5'-TGC TCT CGC GAG GTC GCT TCT CTT-3'), and Enterococcaceae[4] (6-FAM-labeled probes: 5'-GAA AGC GCC TTT CAC TCT TAT GC-3').

FISH was conducted as previously reported with modifications[5]. Briefly, biofilms on PE films were firstly fixed with 4% paraformaldehyde overnight at 4°C. After rinsing in PBS, samples were permeabilized with 1 mg/ml lysozyme (Sigma-Aldrich) for 10 min at room temperature, followed by PBS rinses. Samples were then dehydrated in ethanol series (50%, 70%, 100%) for 3 min each. Hybridization was performed at 46°C for 3 h with a buffer containing 0.9 M NaCl, 20 mM Tris/HCl (pH 7.2), 30% formamide, 0.01% SDS, and fluorochrome-labeled probes (5 ng/μl). Post-hybridization, PE samples were rinsed in washing buffer (20 mM Tris/HCl, 5 mM EDTA, 102 mM NaCl) at 48°C, incubated for 15 min, and rinsed with ice-cold $dH_2O$. FISH samples were imaged with CLSM (LSM 800, Zeiss) and recorded Z-stacks for 3D visualization with 1 μm steps. Stacked images were merged and analyzed using ImageJ software as previously reported[6].

**Text S5. Transcriptomic analyses.** The SynCom and the Ps monocultures were incubated in PE plastics in LCFBM medium for a week, which were exploited as experimental groups. In contrast, populations and single strains grown in LB medium were used as a control group.

Briefly, total RNAs were extracted using TRIzol reagent (Invitrogen, USA) and DNA contamination was removed using MEGA clear Kit (Life technologies, USA). Only high-quality RNA sample was used to construct sequencing library. Ribosome RNA depletion was performed using the RiboCop rRNA Depletion Kit (Lexogen, USA), followed by fragmentation of mRNA into 200 nt fragments. Double-stranded cDNA was synthesized with random hexamer primers (Illumina), incorporating dUTP in the second strand. RNA-seq libraries were prepared using the Illumina Stranded mRNA Prep kit and sequenced on an Illumina Novaseq 6000. Low-quality reads, reads with >10% N bases, and adaptor sequences were removed from the data. For Ps monocultures, high quality reads in each sample were mapped to the reference genome (GCF_019702405.1) of using Bowtie2. Gene and transcript abundances from RNA-Seq data are quantified using RSEM, which employs the Expectation Maximization (EM) algorithm to estimate abundances, accounting for paired-end reads, fragment lengths, and quality scores. Expression levels are calculated using FPKM (fragments per kilobase per million mapped reads) and TPM (transcripts per million), which normalize for gene length and sequencing differences, allowing for direct comparison of gene expression across samples. Differentially expressed genes (DEGs) were identified using the edgeR, DESeq2, or DESeq packages for each dataset and alignment/quantification protocol. As for functional characterization of DEGs, KOBAS and Goatools were used to identify significantly enriched KEGG functions and GO terms, respectively.


1. Bombelli, P., Howe, C. J. & Bertocchini, F. Polyethylene bio-degradation by caterpillars of the wax moth Galleria mellonella. *Curr. Biol.* **27**, R292–R293 (2017).
2. Gunasekera, T. S., Dorsch, M. R., Slade, M. B. & Veal, D. A. Specific detection of Pseudomonas spp. in milk by fluorescence in situ hybridization using ribosomal RNA directed probes. *J. Appl. Microbiol.* **94**, 936–945 (2003).
3. Ootsubo, M. *et al.* Oligonucleotide probe for detecting Enterobacteriaceae by in situ hybridization. *J. Appl. Microbiol.* **93**, 60–68 (2002).
4. Chávez de Paz, L. Development of a Multispecies Biofilm Community by Four Root Canal Bacteria. *J. Endod.* **38**, 318–23 (2012).
5. Selective enumeration of viable Enterobacteriaceae and Pseudomonas spp. in milk within 7 h by multicolor fluorescence in situ hybridization following microcolony formation. *J. Biosci. Bioeng.* **113**, 746–750 (2012).
6. Lin, Y., Xu, X., Maróti, G., Strube, M. L. & Kovács, Á. T. Adaptation and phenotypic diversification of Bacillus thuringiensis biofilm are accompanied by fuzzy spreader morphotypes. *Npj Biofilms Microbiomes* **8**, (2022).

**Supplementary Figures**

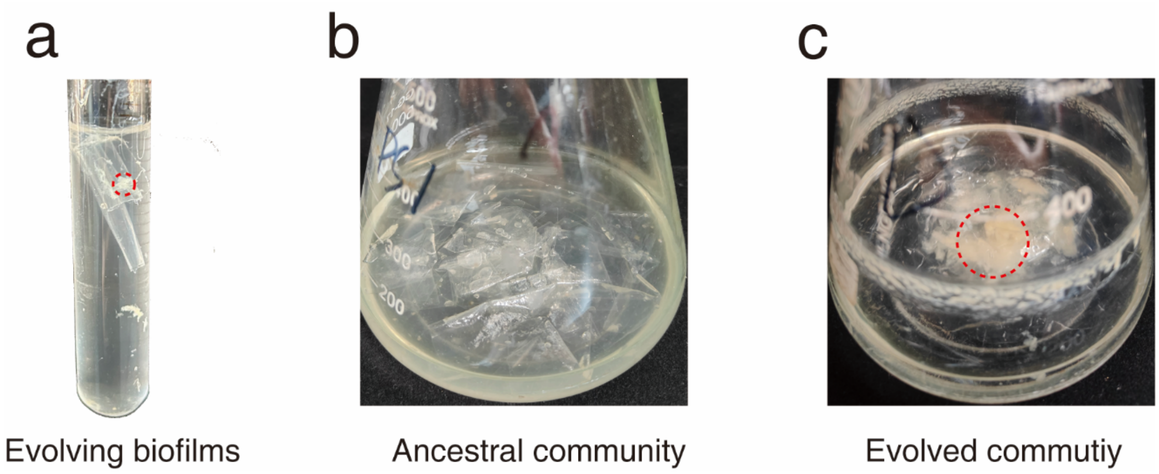


Figure S1. Biofilm formation of evolved populations. (a) The evolved microbial community exhibits a significantly enhanced ability to form cell aggregates. (b) The biofilm formation of the evolved microbial community after one month of long-term culture is compared with that of the ancestral community.

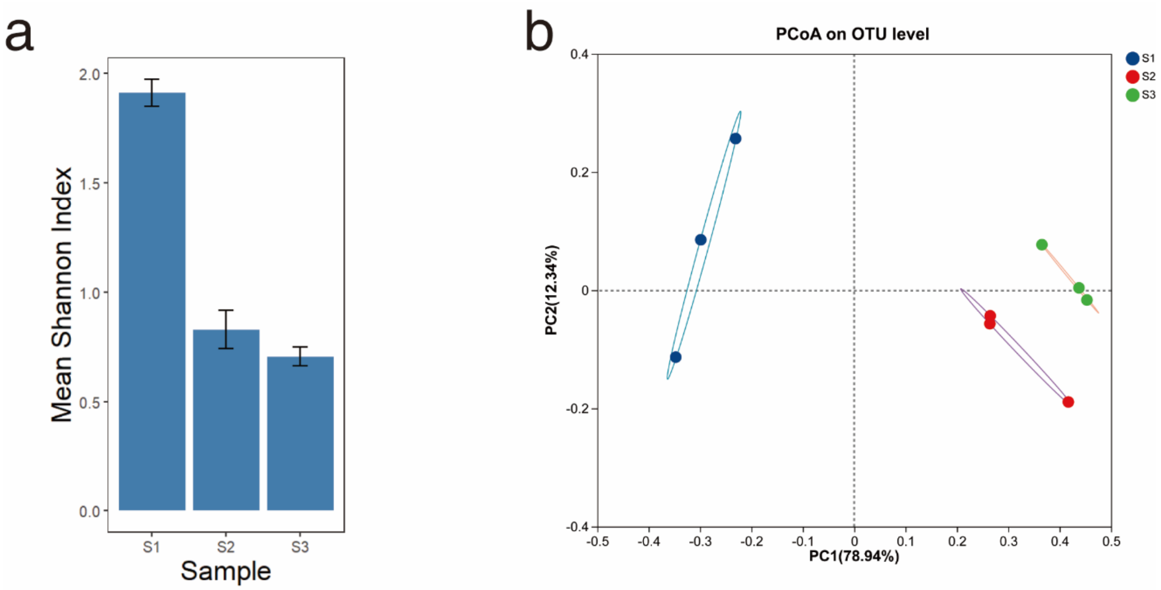


Figure S2. Analysis of microbial community diversity of population 6 during evolution. Three replicates were taken for each time points. (a) Shannon indices at three time points during evolution. (b) PCoA analysis of the evolved microbial community.

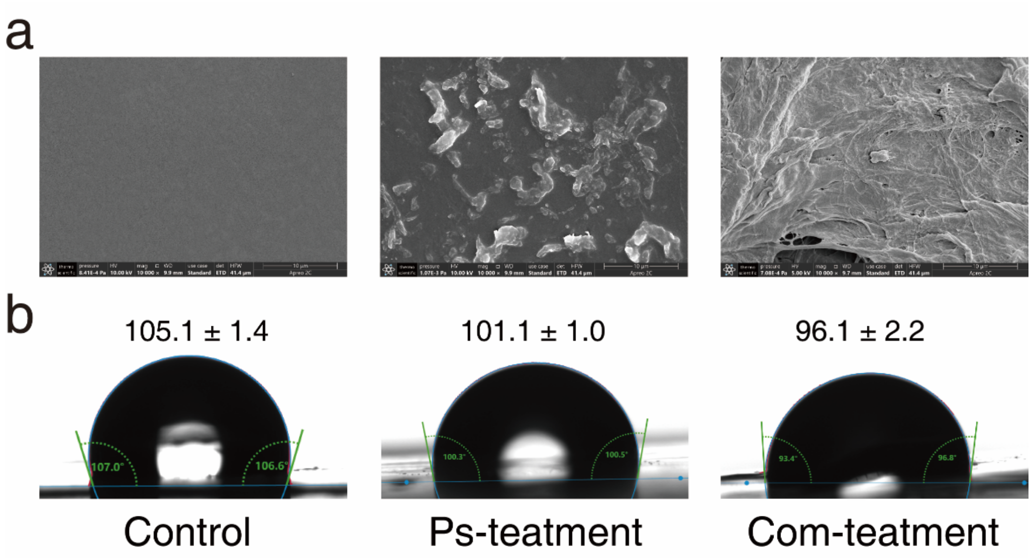


Figure S3. Degradation evidence of Ps and the SynCom incubated with PE for 30 days. (a) PE surface changes after 30 days of incubation with Ps and the SynCom. (b) Water contact angle experiment on PE plastic treated with bacteria.

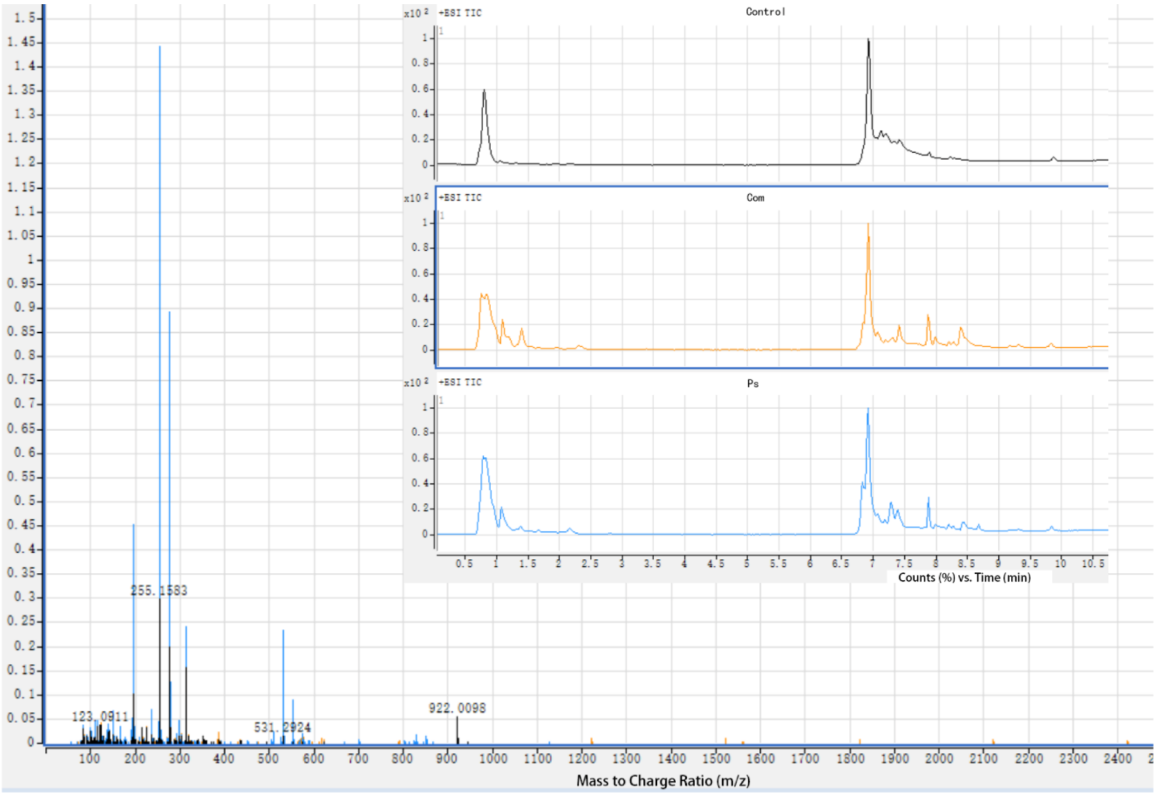


Figure S4. HPLC-MS spectrum of products released from the PE films treated with Ps, SynCom or medium as control for a month.

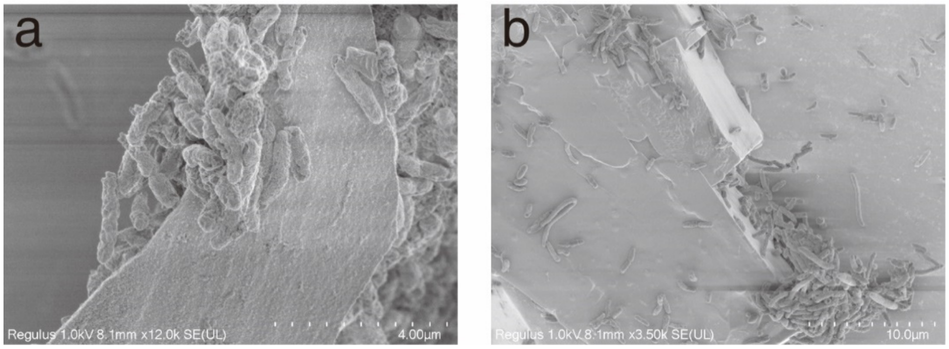


Figure S5. SEM analysis of surface bacteria during PE plastic degradation. (a) SEM images of Ps on PE surface. (b) SEM images of the SynCom on PE surface.

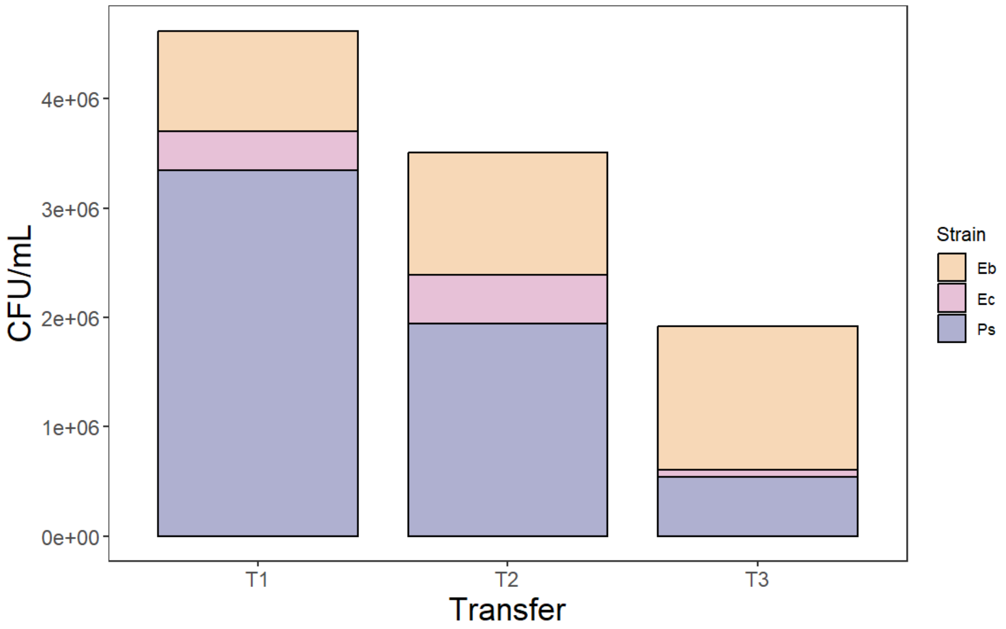


Figure S6. Biofilm formation on PE plastic after the disruption of biofilm structure of microbial community.

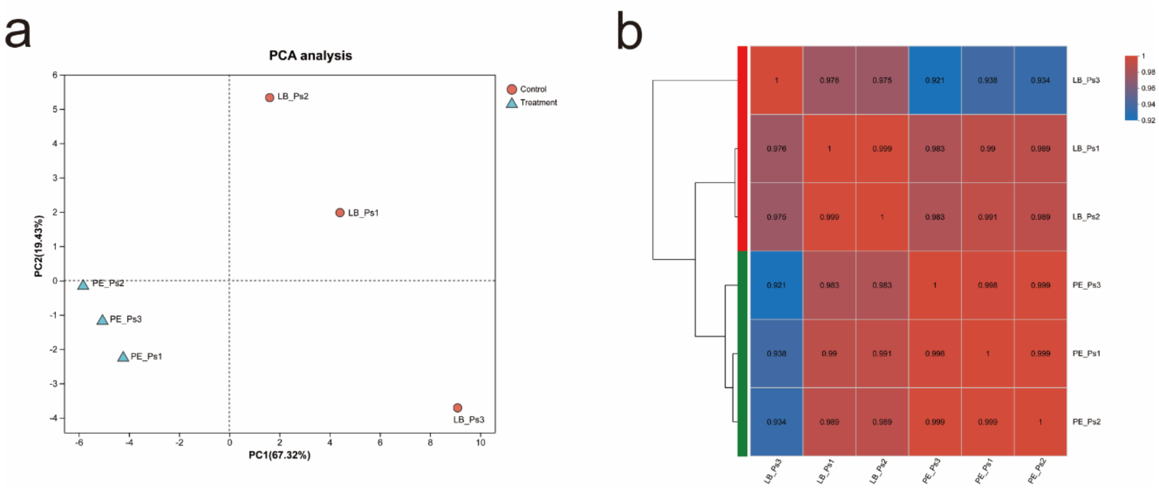


Figure S7. Expression level analysis between samples. (a) PCA analysis between samples in PE and LB treatment. (b) Correlation analysis of expressed genes between samples in PE and LB treatment.

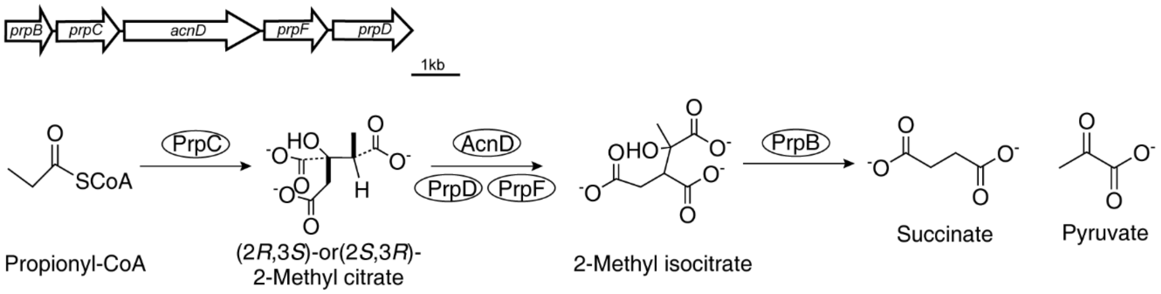


Figure S8. *prp* operon in *Pseudomonas spp*. and propionyl-CoA metabolism pathways.

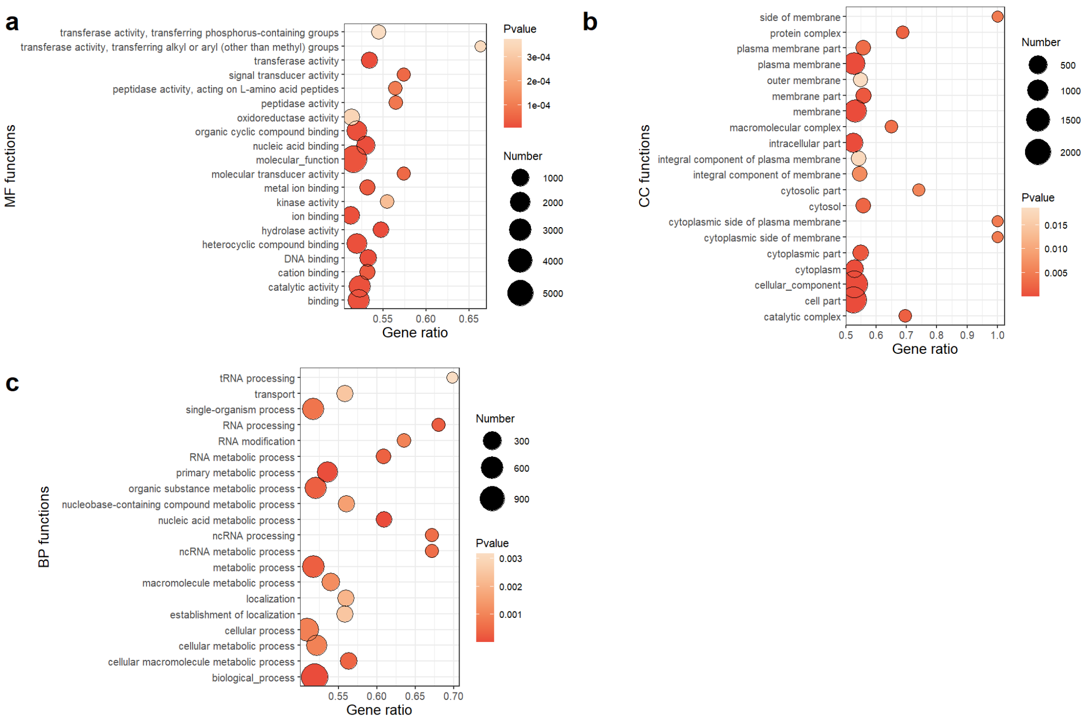


Figure S9. GO enrichment analysis of significantly upregulated genes in the SynCom during cultivation in PE compared to LB medium. (a) Molecular functions. (b) Cellular components. (c) Biological processes.
